# Assessing combinatorial diversity of aureochrome bZIPs through genome-wide screening

**DOI:** 10.1101/2022.06.05.494858

**Authors:** Madhurima Khamaru, Deep Nath, Devrani Mitra, Soumen Roy

## Abstract

Aureochromes are unique blue light-responsive LOV (Light Oxygen Voltage) photoreceptors cum basic leucine zipper (bZIP) transcription factors (TFs), present exclusively in photosynthetic marine stramenopiles. Considering the availability of the complete genome sequence, this study focuses particularly on aureochromes from *Ectocaupus siliculosus*. Aureochromes mediate light-regulated developmental responses in this brown photosynthetic algae. Both the LOV sensor and the bZIP effector shows sequence-structure conservation. The LOV+bZIP modules of aureochrome homologs/ paralogs are not only structurally similar but also show an identical oligomeric state -- preferably dimeric. Aureochromes execute diverse cellular responses in different photosynthetic stramenopiles-- though their activities can vary even within a given algal species. Besides a heterogeneous linker connecting the sensor-effector and a flexible N-terminal region, the sequence composition of both the domains is vital. Therefore, it is important to understand whether aureochromes select dimerization partners from the same family or interact with other bZIPs as well. To regulate multifarious biological activities, it is possible that aureochromes activate the global TF interaction network. Following homo/heterodimer modeling, we address the compatibility of dimerization partners by screening through heptad repeats. We evaluate the dimer interface area in terms of gain in solvation energy as well as the number of hydrogen bonds/salt bridge interactions. We further explore the relative stability of these structures from a graph-theoretic perspective through well-studied measures such as the energy of the graph and average participation coefficient. Furthermore, we also conduct an information-theoretic analysis using network information centrality and Kullback-Leibler divergence. We find that all our investigations into the relative stability of these dimers using diverse methods from bioinformatics, network science, and, information theory are in harmonious agreement. Our approach and findings should facilitate the design of experiments.

## Introduction

The intrinsic cellular processes governing life are regulated by transcription factors (TFs). They bridge the gene regulation pattern of cells with diverse extracellular or intracellular signaling cascades (Weidemüller et al., 2021). TFs play a crucial operative part in diverse intricate gene regulatory networks. They can either directly bind to specific DNA sequences like promoters, gene regulatory regions (enhancers/silencers), or, can engage in protein–protein interactions with other TFs. TFs may undergo oligomerization (dimers or multimers), facilitated through hydrogen bonding and salt bridge formation to bind with elements in the target DNA. Coordinated interaction between TFs, regulatory proteins and co-factors at the enhancer/promoter site drives differential expression of multiple genes in a cell specific manner. Such spatio-temporally precise gene expression events ultimately guide the growth and developmental response of living organisms.

Among several well known subfamilies of TFs, basic leucine zippers (bZIPs) demonstrate high conservation of their sequence in the basic region as well as their structure. bZIPs bind with specificDNA substrates essentially as dimers (Deppmann et al., 2004a)(Grigoryan & Keating, 2006). The bipartite α-helical structure of the dimer bZIPs forms a ‘Y’ shaped complex with the DNA(Bader & Vogt, 2006b). In fact, dimerization among compatible bZIP family members has resulted into a genome wide dimer mediated regulatory network of bZIPs (Li et al., 2022). And, it is still expanding with the gene duplication of the different bZIP sequences. Aureochromes are considered to be a rare amalgamation of a photoreceptor and a TF, with bZIPs as an effector (Khamaru et al., 2022). A sensory TF like aureochrome can activate the global TF interaction network not only by interacting with the DNA but also by finding compatible monomeric bZIP partners. This gives rise to potential combinatorial diversity, allowing them to choose from a wide range of substrate DNA repertoire and execute diverse biological activities. While the homodimerization of aureochromes from several organisms is supported by both *in vitro/in vivo* experiments, heterodimerization between aureochrome bZIP monomers is reported only in *Phaeodactylum sp* (Banerjee et al., 2016). This study therefore focusses on homo/heterodimerization of aureochrome bZIPs as well as other other bZIPs through genome wide screening. We choose photosynthetic brown algae, *Ectocarpus siliculosus* as model organism (Cock et al., 2010).

Various studies on the homo/heterodimerization potential of human bZIPs have revealed several critical residues from the leucine-zipper sequence as well as specific inter-helical interactions that aid in dimer specificity and stabilization (C. Vinson et al., 2002). For example, the residues present at the a, g, e positions of each heptad consisting of seven amino acid residues (a-b-c-d-e-f-g), determine the preference for the dimerization partners of bZIPs(C. R. Vinson et al., 1993)(Krylov et al., 1994). The presence of an asparagine residue at ‘a’ increases the chance of homodimerization, whereas a charged residue at ‘a’ would inhibit homodimer formation. Lysine substitution in this position would however lead towards heterodimerization (C. Vinson et al., 2002). Quite obviously, if identically charged residues occupy g-e’ positions (where e’ is located in the adjacent heptad, five residues downstream), the formation of homodimer is unlikely. A well-known bZIP TF, Fos that contains glutamate in such positions is unable to form homodimer (Deppmann et al., 2004b).

The dimerization property of any bZIP is of immense importance for its substrate DNA specificity and gene regulatory activity. But the dimerization properties of aureochrome bZIPs, especially across genome are unexplored till date. In respect to domain topology, aureochromes generally possess an N-terminal bZIP as an effector and C-terminal LOV as a sensor — yet aureochrome subtypes within an organism or from multiple organisms, execute diverse range of biological activities. This includes photomorphogenesis (Takahashi et al., 2007) (Deng et al., 2014), lipid accumulation (Huang et al., 2014), cell cycle progression (Mann et al., 2020), photo acclimation (Kroth et al., 2017) etc. Therefore, the fundamental question is despite structural identity, how such differences in activities ariseti There could be multiple possible factors including heterogeneity of linker connecting effector and sensor domain, uncharacterized N-terminal region of variable length and sequence, subtle differences in amino acid sequence composition of the sensor as well as effector that influences photoabsorption and leucine zipper dimerization/stability respectively. This paper particularly deals with the latter aspect. We investigate the potential of aureochrome bZIPs towards partner selection, necessary for dimerization, ultimately manifested in different substrate DNA binding activity and respective physiological processes. Screening through *Ectocarpus* genome, we create several bZIP combinations possible for aureochromes and investigate their dimerization properties behind huge combinatorial diversity. Auer protein dimer modeling, we analyze the compatibility/stability of dimers using tools from structural bioinformatics including molecular dynamics (MD) simulation. We further construct networks of these modeled structures (Grewal, Roy, 2015) (Deb, Grewal et al., 2020). We study the stability of these structures by evaluating the participation coefficient (Di Paola, Platania, et al., 2015) and energy of these networks (Di Paola, Mei, et al., 2015). In addition, we also conduct investigations using information theoretic measures like information centrality and Kullback-Leibler divergence. Finally, we test these mathematical predictions by interface analysis and signature Molecular Dynamics (MD) simulations on selected structures. We find that our findings from all these agreement are in harmonious agreement with each other.

## Materials and Methods

1. **Sequence retrieval**-Considering the availability of whole genome sequence of *Ectocarpus siliculosus* (Cock, Sterck et al., 2010), we searched for the bZIPs present in it. From the Uniprot knowledgebase, the sequences of all bZIPs including those from aureochromes are retrieved.
2. **Dimer modeling**-All homo and heterodimers are modeled using SWISS-MODEL (Waterhouse et al., 2018). Following Ramachandran plot analysis, these modeled dimers are then energy minimized using Chimera. Ramachandran statistics are verified again by Procheck.
3. **Interface analysis of dimers**-All the dimer models are then subjected to PDBePISA analysis to calculate the interface area, gain in solvation free upon dimerization, as well as the number of hydrogen bonds and salt bridges formed.
4. **Construction of the graph-** Every residue of the protein is considered to be a node in the network or graph, denoted henceforth by G. For all residues, we evaluate the distance between every pair of atoms — where each atom belongs to a different residue. If the minimum distance between a pair of such atoms is lesser or equal to 4.5Å, then the residues to which these atoms belong, are connected by an edge in the graph (Grewal, Roy, 2015). Such graphs have also been constructed to study Aureochromes (Grewal, Mitra, 2015). The network thus obtained is thence subjected to further analyses. We denote the number nodes in the network by N.
5. **Energy of a graph-** The energy, E_G_, of a graph, G, is the sum of absolute values (modulus) of the eigenvalues of the adjacency matrix of G (Di Paola, Platania, et al., 2015). In this paper, two monomers have been used to construct every dimer. Thus, we have three graph structures — those of the two monomers and the resultant dimer. Let us denote the energy of the dimer as E_D_ and of the monomers as E_M1_ and E_M2_. Energy difference upon dimerization, ΔE = E_D_ - (E_M1_ + E_M2_). The normalized form of ΔE, also known as specific difference of energy is ΔE_SP_ = ΔE/N_D_; where, N_D_ is the total number of residues in the dimer. Higher graph energy indicates higher stability of the dimer. Further, the interface of the dimer plays an important role in its stability (Di Paola, Mei, et al., 2015). The graph energy accords a tool from the perspective of topological analyses of networks to study the stability of a protein. It helps in identifying the role of inter-subunit surface in protein-folding and offers fresh and helpful insight over traditional measures such as Gibbs free energy, which have been extensively studied in systems not amenable to graphical analyses (Di Paola, Mei, et al., 2015).
6. **Participation Coefficient-** Participation coefficient, P_i_, of node, i, in a graph is defined as P_i_=1-(k_si_/k_i_)^2^ (Di Paola, Platania, et al., 2015). Here, each monomer of the dimer is considered as a cluster and k_si_ indicates the fraction of total nodes in the cluster to which node, i, belongs — which are connected to i. k_i_ represents the fraction of total nodes in the entire network to which node, i, is connected. P_i_ signifies the strength of connection of the i^th^ node with others nodes in its cluster. It has been reported that P_i_ is zero for residues close to an allosteric site. It has also been shown that nodes with a high value of P_i_ contribute significantly towards the stability of that protein (Di Paola, Platania, et al., 2015). P_i_ has been calculated to understand inter-monomeric connections. Further, the average value of P_i_, P_av_ = (∑_i_ P_i_) / N_D_, is calculated to understand the overall stability of the dimer.
7. **Kullback-Leibler divergence-** Kullback-Leibler (KL) divergence or relative entropy aims to measure the manner in which two distributions differ from each other (Kullback & Leibler, 1951). For two discrete distributions, A and B, the relative entropy of A to B is determined as, 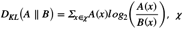, being the space in which both A and B are defined. Henceforth, *D*_*KL*_(*A* ‖ *B*) would be denoted simply by *D*_*KL*_. Here, A denotes the degree distribution of the dimer. B denotes the degree distribution of the hypothetical graph created by a contrived merger of both monomers.
8. **Information Centrality-** In its original form, information centrality, I_i,_ of node, i, essentially measures the harmonic mean length of paths ending at node i. I_i_ is lower if node i has many short paths connecting other vertices to it (Stephenson & Zelen, 1989). This information measure has a clear interpretation in electrical network theory. It can be thought of as the effective resistance between two nodes in G, when a resistor is associated with every edge of G (Brandes & Fleischer, 2005). Herein, we average the value of I_i_ over all nodes in G. We denote average information centralities of dimer and monomers as I_D_, I_M1_ and I_M2_ respectively. We are interested in relative change of I_D_, expressed as I_r_ = [(I_M1_ + I_M2_) - I_D_] / (I_M1_ + I_M2_). Values of I_r_ appear in Table 1. Values of I_D_, I_M1_, and, I_M2_ appear in Supplementary Information.

**Table 1:**
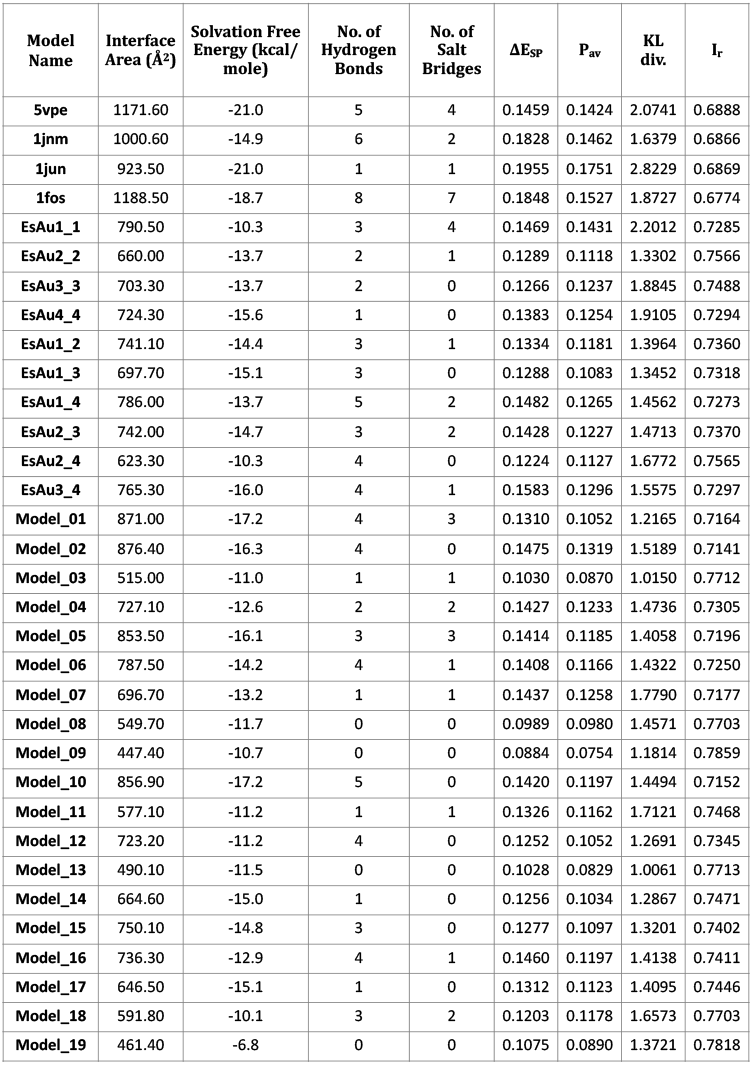
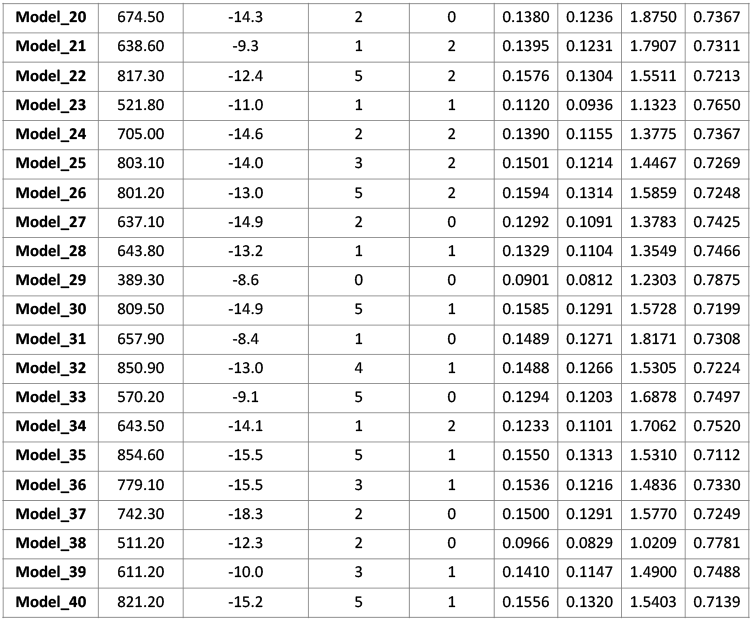
Values of specific difference of energy (ΔE_SP_), average participation coefficient (P_av_), Kullback-Leibler divergence (KL), and, relative information centrality (I_r_) for all possible dimers. Esi_0219_0040 in complex with EsAureo1, EsAureo2, EsAureo3 and EsAureo4 represents Model_1, Model_11, Model_21, Model_31 respectively. Esi_0113_0081 in complex with EsAureo1, EsAureo2, EsAureo3 and EsAureo4 represents Model_2, Model_12, Model_22, Model_32 respectively. Esi_0039_0117 in complex with EsAureo1, EsAureo2, EsAureo3 and EsAureo4 represents Model_3, Model_13, Model_23, Model_33 respectively. Esi_0199_0057 in complex with EsAureo1, EsAureo2, EsAureo3 and EsAureo4 represents Model_4, Model_14, Model_24, Model_34 respectively. Esi_0145_0005 in complex with EsAureo1, EsAureo2, EsAureo3 and EsAureo4 represents Model_5, Model_15, Model_25, Model_35 respectively. Esi_0065_0095 in complex with EsAureo1, EsAureo2, EsAureo3 and EsAureo4 represents Model_6, Model_16, Model_26, Model_36 respectively. Esi_0211_0027 in complex with EsAureo1, EsAureo2, EsAureo3 and EsAureo4 represents Model_7, Model_17, Model_27, Model_37 respectively. Esi_0017_0151 in complex with EsAureo1, EsAureo2, EsAureo3 and EsAureo4 represents Model_8, Model_18, Model_28, Model_38 respectively. Esi_0060_0098 in complex with EsAureo1, EsAureo2, EsAureo3 and EsAureo4 represents Model_9, Model_19, Model_29, Model_39 respectively. Esi_0035_0131 in complex with EsAureo1, EsAureo2, EsAureo3 and EsAureo4 represents Model_10, Model_20, Model_30, Model_40 respectively.
9. **Molecular Dynamics Simulation**-The MD simulations are performed using GROMACS version 2021.3 & version 2022. In each case, charm27 all-atom force field is employed with SPC water model and cubic simulation box. The solvated protein is then neutralized with sodium and chloride ions. Steepest descent method (50000 steps) is next employed to energy minimize the neutralized system. The energy minimized system is then subjected to NVT equilibration simulation for 100 ps such that the system aiains a temperature of 300K. This is followed by 100 ps NPT equilibration at 1 atm pressure maintaining identical temperature. The temperature and pressure are maintained by Berendsen thermostat and Parrinello-Raheman pressure. For calculating long range electrostatic forces, Particle-Mesh Ewald (PME) is applied. Each NPT equilibrated system is then subjected to 20ns production run with 2fs time step without any restraint. The trajectory analysis of the MD simulation is performed using the GROMACS command line tools. Root mean square deviation (RMSD) and root mean square fluctuation (RMSF) of the protein backbone are then analyzed. The status of interchain hydrogen bonds within the dimer in the protein-protein complex is also calculated.

| **Model Name** | **Interface Area (Å<sup>2</sup>)** | **Solvation Free Energy (kcal/mole)** | **No. of Hydrogen Bonds** | **No. of Salt Bridges** | **ΔE<sub>SP</sub>** | **P<sub>av</sub>** | **KL div.** | **I<sub>r</sub>** |
| --- | --- | --- | --- | --- | --- | --- | --- | --- |
| **5vpe** | 1171.60 | -21.0 | 5 | 4 | 0.1459 | 0.1424 | 2.0741 | 0.6888 |
| **1jnm** | 1000.60 | -14.9 | 6 | 2 | 0.1828 | 0.1462 | 1.6379 | 0.6866 |
| **1jun** | 923.50 | -21.0 | 1 | 1 | 0.1955 | 0.1751 | 2.8229 | 0.6869 |
| **1fos** | 1188.50 | -18.7 | 8 | 7 | 0.1848 | 0.1527 | 1.8727 | 0.6774 |
| **EsAu1_1** | 790.50 | -10.3 | 3 | 4 | 0.1469 | 0.1431 | 2.2012 | 0.7285 |
| **EsAu2_2** | 660.00 | -13.7 | 2 | 1 | 0.1289 | 0.1118 | 1.3302 | 0.7566 |
| **EsAu3_3** | 703.30 | -13.7 | 2 | 0 | 0.1266 | 0.1237 | 1.8845 | 0.7488 |
| **EsAu4_4** | 724.30 | -15.6 | 1 | 0 | 0.1383 | 0.1254 | 1.9105 | 0.7294 |
| **EsAu1_2** | 741.10 | -14.4 | 3 | 1 | 0.1334 | 0.1181 | 1.3964 | 0.7360 |
| **EsAu1_3** | 697.70 | -15.1 | 3 | 0 | 0.1288 | 0.1083 | 1.3452 | 0.7318 |
| **EsAu1_4** | 786.00 | -13.7 | 5 | 2 | 0.1482 | 0.1265 | 1.4562 | 0.7273 |
| **EsAu2_3** | 742.00 | -14.7 | 3 | 2 | 0.1428 | 0.1227 | 1.4713 | 0.7370 |
| **EsAu2_4** | 623.30 | -10.3 | 4 | 0 | 0.1224 | 0.1127 | 1.6772 | 0.7565 |
| **EsAu3_4** | 765.30 | -16.0 | 4 | 1 | 0.1583 | 0.1296 | 1.5575 | 0.7297 |
| **Model_01** | 871.00 | -17.2 | 4 | 3 | 0.1310 | 0.1052 | 1.2165 | 0.7164 |
| **Model_02** | 876.40 | -16.3 | 4 | 0 | 0.1475 | 0.1319 | 1.5189 | 0.7141 |
| **Model_03** | 515.00 | -11.0 | 1 | 1 | 0.1030 | 0.0870 | 1.0150 | 0.7712 |
| **Model_04** | 727.10 | -12.6 | 2 | 2 | 0.1427 | 0.1233 | 1.4736 | 0.7305 |
| **Model_05** | 853.50 | -16.1 | 3 | 3 | 0.1414 | 0.1185 | 1.4058 | 0.7196 |
| **Model_06** | 787.50 | -14.2 | 4 | 1 | 0.1408 | 0.1166 | 1.4322 | 0.7250 |
| **Model_07** | 696.70 | -13.2 | 1 | 1 | 0.1437 | 0.1258 | 1.7790 | 0.7177 |
| **Model_08** | 549.70 | -11.7 | 0 | 0 | 0.0989 | 0.0980 | 1.4571 | 0.7703 |
| **Model_09** | 447.40 | -10.7 | 0 | 0 | 0.0884 | 0.0754 | 1.1814 | 0.7859 |
| **Model_10** | 856.90 | -17.2 | 5 | 0 | 0.1420 | 0.1197 | 1.4494 | 0.7152 |
| **Model_11** | 577.10 | -11.2 | 1 | 1 | 0.1326 | 0.1162 | 1.7121 | 0.7468 |
| **Model_12** | 723.20 | -11.2 | 4 | 0 | 0.1252 | 0.1052 | 1.2691 | 0.7345 |
| **Model_13** | 490.10 | -11.5 | 0 | 0 | 0.1028 | 0.0829 | 1.0061 | 0.7713 |
| **Model_14** | 664.60 | -15.0 | 1 | 0 | 0.1256 | 0.1034 | 1.2867 | 0.7471 |
| **Model_15** | 750.10 | -14.8 | 3 | 0 | 0.1277 | 0.1097 | 1.3201 | 0.7402 |
| **Model_16** | 736.30 | -12.9 | 4 | 1 | 0.1460 | 0.1197 | 1.4138 | 0.7411 |
| **Model_17** | 646.50 | -15.1 | 1 | 0 | 0.1312 | 0.1123 | 1.4095 | 0.7446 |
| **Model_18** | 591.80 | -10.1 | 3 | 2 | 0.1203 | 0.1178 | 1.6573 | 0.7703 |
| **Model_19** | 461.40 | -6.8 | 0 | 0 | 0.1075 | 0.0890 | 1.3721 | 0.7818 |
| Model_20 | 674.50 | -14.3 | 2 | 0 | 0.1380 | 0.1236 | 1.8750 | 0.7367 |
| Model_21 | 638.60 | -9.3 | 1 | 2 | 0.1395 | 0.1231 | 1.7907 | 0.7311 |
| Model_22 | 817.30 | -12.4 | 5 | 2 | 0.1576 | 0.1304 | 1.5511 | 0.7213 |
| Model_23 | 521.80 | -11.0 | 1 | 1 | 0.1120 | 0.0936 | 1.1323 | 0.7650 |
| Model_24 | 705.00 | -14.6 | 2 | 2 | 0.1390 | 0.1155 | 1.3775 | 0.7367 |
| Model_25 | 803.10 | -14.0 | 3 | 2 | 0.1501 | 0.1214 | 1.4467 | 0.7269 |
| Model_26 | 801.20 | -13.0 | 5 | 2 | 0.1594 | 0.1314 | 1.5859 | 0.7248 |
| Model_27 | 637.10 | -14.9 | 2 | 0 | 0.1292 | 0.1091 | 1.3783 | 0.7425 |
| Model_28 | 643.80 | -13.2 | 1 | 1 | 0.1329 | 0.1104 | 1.3549 | 0.7466 |
| Model_29 | 389.30 | -8.6 | 0 | 0 | 0.0901 | 0.0812 | 1.2303 | 0.7875 |
| Model_30 | 809.50 | -14.9 | 5 | 1 | 0.1585 | 0.1291 | 1.5728 | 0.7199 |
| Model_31 | 657.90 | -8.4 | 1 | 0 | 0.1489 | 0.1271 | 1.8171 | 0.7308 |
| Model_32 | 850.90 | -13.0 | 4 | 1 | 0.1488 | 0.1266 | 1.5305 | 0.7224 |
| Model_33 | 570.20 | -9.1 | 5 | 0 | 0.1294 | 0.1203 | 1.6878 | 0.7497 |
| Model_34 | 643.50 | -14.1 | 1 | 2 | 0.1233 | 0.1101 | 1.7062 | 0.7520 |
| Model_35 | 854.60 | -15.5 | 5 | 1 | 0.1550 | 0.1313 | 1.5310 | 0.7112 |
| Model_36 | 779.10 | -15.5 | 3 | 1 | 0.1536 | 0.1216 | 1.4836 | 0.7330 |
| Model_37 | 742.30 | -18.3 | 2 | 0 | 0.1500 | 0.1291 | 1.5770 | 0.7249 |
| Model_38 | 511.20 | -12.3 | 2 | 0 | 0.0966 | 0.0829 | 1.0209 | 0.7781 |
| Model_39 | 611.20 | -10.0 | 3 | 1 | 0.1410 | 0.1147 | 1.4900 | 0.7488 |
| Model_40 | 821.20 | -15.2 | 5 | 1 | 0.1556 | 0.1320 | 1.5403 | 0.7139 |

## Results

### Sequence retrieval and dimer modeling

Dimerization among suitable bZIP partners is essential for the selection of diverse substrate DNA and the regulation of multiple biological activities. This study focuses on unique photoreceptor-TF, Aureochromes from photosynthetic brown alga, *Ectocarpus siliculosus*. The whole genome sequence of *Ectocarpus* (Cock et al. 2010) reveals the presence of five aureochrome subtypes, in which aureochrome5 contains only the LOV sensor. Therefore, we consider only four subtypes of aureochromes (1-4) and investigate their compatibility not only as homodimers but also as inter-subtype heterodimers. We further consider remaining 10 bZIPs, deciphered in *Ectocarpus* genome sequence (Cock et al. 2010) and investigate their compatibility with aureochrome bZIPs via heterodimerization. In absence of experimental 3D structures, we conduct monomer/dimer modeling using SWISS-MODEL server or by Pymol. We generate 4 homodimers, 6 inter-subtype heterodimers and 40 heterodimers between aureochromes and 10 other bZIPs. The potential compatibility among all bZIPs, investigated in this study is presented in **Figure 1**. The respective sequence information is collected from Uniprot database prior to modeling. Auer validation through Ramachandran plot analysis and energy minimization using Chimera, we characterize the dimer interface in terms of amino acid residue composition, interface area, gain in energy of solvation, bonding interactions, graph-theoretic and information-theoretic analyses.

**Figure 1:**
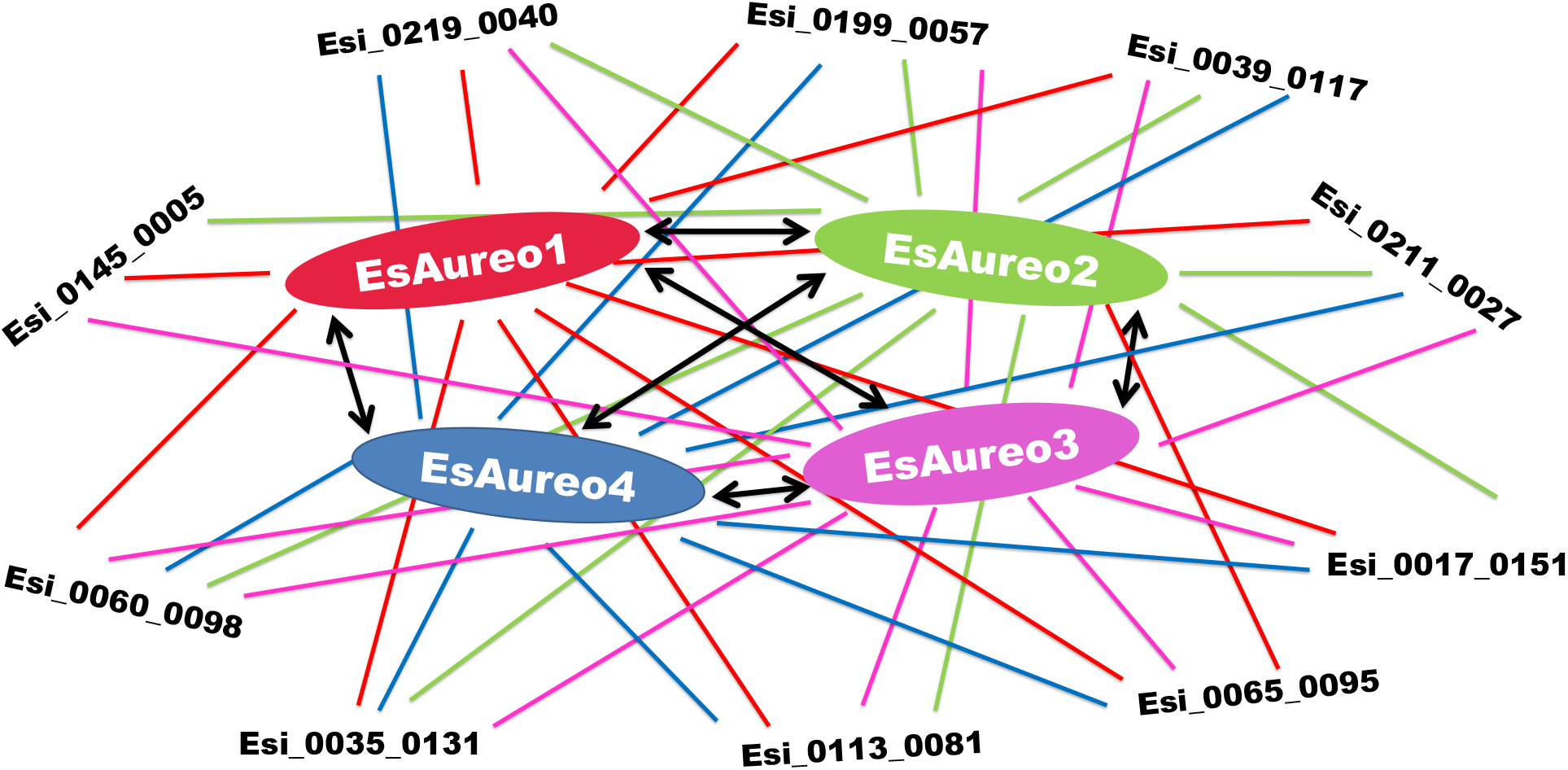
Possibilities of homo/hetero-dimerization involving *E. siliculosus a*ureochromes and other basic leucine zippers

### Characterization of dimer interface

To investigate the reliability of the modeled dimers and understand the compatibility the energy minimized structures are subjected to PDBePISA (Protein interfaces surfaces and assemblies) (Krissinel and Henrick, 2007). The solvation energy (ΔG), interface area, number of hydrogen bonding and salt bridges interactions are studied thoroughly. The results are presented in **Table 1**. Detailed information is available in SI [**Table SI_1**]. Besides modeled dimer structures, we also include crystal structures of several bZIP homo/heterodimers for effective comparison. Negative values of gain in solvation free energy for all modeled dimers indicate potential protein-protein complex formation between the bZIP partners. While the interface area in experimentally determined bZIP complexes ranges from about 900Å^2^ to 1200 Å^2 —^ in modeled dimer structures, it ranges from about 400Å^2^ to 900 Å^2^. Perhaps it is worth mentioning that except sequences from bZIP crystal structures, we generally considered about 50 residues for modeling. However, in few cases e.g., Esi_0039_0117 or Esi_0060_0098, 9 to 10 residues at the C-terminal could not be modeled, possibly due to sequence heterogeneity and no homology with any of the nearly 50 possible template sequences. The missing C-terminal residues in such cases do not significantly disturb heptad repeats, although they might have contributed towards a lower interface area. In fully modeled dimers, the number of salt bridges and hydrogen bonding interactions is comparable with that of experimentally obtained bZIP dimers. Few dimers like model_8, model_9, model_13, model_19, and, model_29 do not show a single hydrogen bonding/salt bridge interaction. Though model_9, model_19, and, model_29 contain one polypeptide chain from Esi_0060_0098 and model_13 contains one Esi_0039_0117 polypeptide, the remaining models constructed by them do show bonding interactions. Therefore, truncation at the C-terminal end might not have influenced the results drastically.

### Prediction of dimer stability following network and information-theoretic approach

The theory of complex networks and information theory has been profoundly useful (Di Paola, Mei, et al., 2015)(Di Paola, Platania, et al., 2015) in understanding the compatibility and stability of protein-protein interaction. Our motivation is to identify structures, which possess relatively higher stability over the others. We therefore calculate the average participation coefficient, P_av_, and specific difference of energy, ΔE_SP,_ of all structures to understand dimer stability (Di Paola, Mei, et al., 2015) (Di Paola, Platania, et al., 2015). In **Table 1**, we observe that the results for P_av_ and ΔE_SP_ demonstrate good mutual agreement. Higher value of ΔE_SP_ of a dimer indicates higher stability. As seen in **Table 2**, values of ΔE_SP_ above a cutoff lead to identification of structures with relatively higher stability. We also identify structures with relatively lower stability.

**Table 2:** List of modeled structures which possess relatively higher stability or lower stability as obtained through diverse measures based on network topology, namely, specific difference of energy (ΔE_SP_), average participation coefficient (P_av_), Kullback-Leibler divergence (KL), and, relative information centrality (I_r_). σ denotes the standard deviation of the respective distribution.

|  | High stability | Low stability |
| --- | --- | --- |
| **$\Delta E_{SP}$** | 1jnm, 1jun, 1fos, EsAu3_4, model_26, model30<br>(1.0s greater than the mean) | model_03, model_08, model_09, model_13, model_19, model_23, model_29, model_38<br>(1.0s less than the mean) |
| **$P_{av}$** | 5vpe, 1jnm, EsAu1_1, 1jun, 1fos<br>(1.0s greater than the mean) | model_03, model_08, model_09, model_13, model_19, model_23, model_29, model_38<br>(1.0s less than the mean) |
| **KL** | 5vpe, EsAu1_1, EsAu3_3, EsAu4_4, model_20, 1jun, 1fos<br>(1.0s greater than the mean) | model_1, model_03, model_09, model_13, model_23, model_38<br>(1.0s less than the mean) |
| **$I_r$** | model_35, 1jnm, 5vpe, 1jun, 1fos<br>(1.0s less than the mean) | model_03, model_08, model_09, model_13, model_18, model_19, model_23, model_29, model_38<br>(1.0s greater than the mean) |

Apart from these two well known measures, we also conduct an information-theoretic investigation of all modeled structures, through KL divergence and network information centrality. KL divergence helps in distinguishing between two probability distributions. In the present context, it serves as a mathematical tool to understand the stability of a dimer. A higher value of KL divergence implies that the structure of the dimer is significantly different from the structure of the monomers. Here, we have considered degree distribution as a signature of the structure. Higher inter-monomer (interchain) connections would also imply a larger dimer interface. As observed in **Table 1** and **2**, the results related to KL divergence agree with the two other well known network measures, P_av_ and ΔE_Sp_. On the other hand, relative information centrality, I_r_, is observed to reflect the overall change of paths in the graph upon dimer formation. For the dimer to be structurally stable, I_D_ has to be high. Higher I_D_ implies a higher number of available paths between two monomers. Therefore, the dimer interface is likely to be higher and the structure is more likely to possess a higher stability. Herein, we are interested in the comparative stability of the dimers. Therefore, instead of I_D_, I_r_ is chosen as a candidate. This choice is suitable to understand the stability of dimer conformation in terms of the available paths between nodes. The variation of I_r_ is observed to be consistent with other measures regarding stability.

While all values of specific difference of energy (ΔE_SP_), average participation coefficient (P_av_), Kullback-Leibler divergence (KL), and, relative information centrality (I_r_) are listed in **Table 1**, detailed information is available in SI [**Table SI_2**] for all candidate dimer models. For each of these measures and for every candidate dimer, we further calculate the standard deviation, σ, of the respective distribution. The implications regarding the stability of a given dimer, as deduced from the value of σ, are summarised in **Table 2**. Based on these measures, we select five stable and four unstable dimers [**Figure 2**] to understand any correlation between interface area, bonding interactions with results from networks and information theory. Model_1 and model_38, which are predicted to be unstable according to KL and Ir values respectively, show sufficient interface area with hydrogen bonding/salt bridge interactions. Therefore, we may conclude that even though the size of the dimer interface or the number of bonding interactions are positively correlated with dimer stability, they are not absolute determinants.

**Figure 2:**
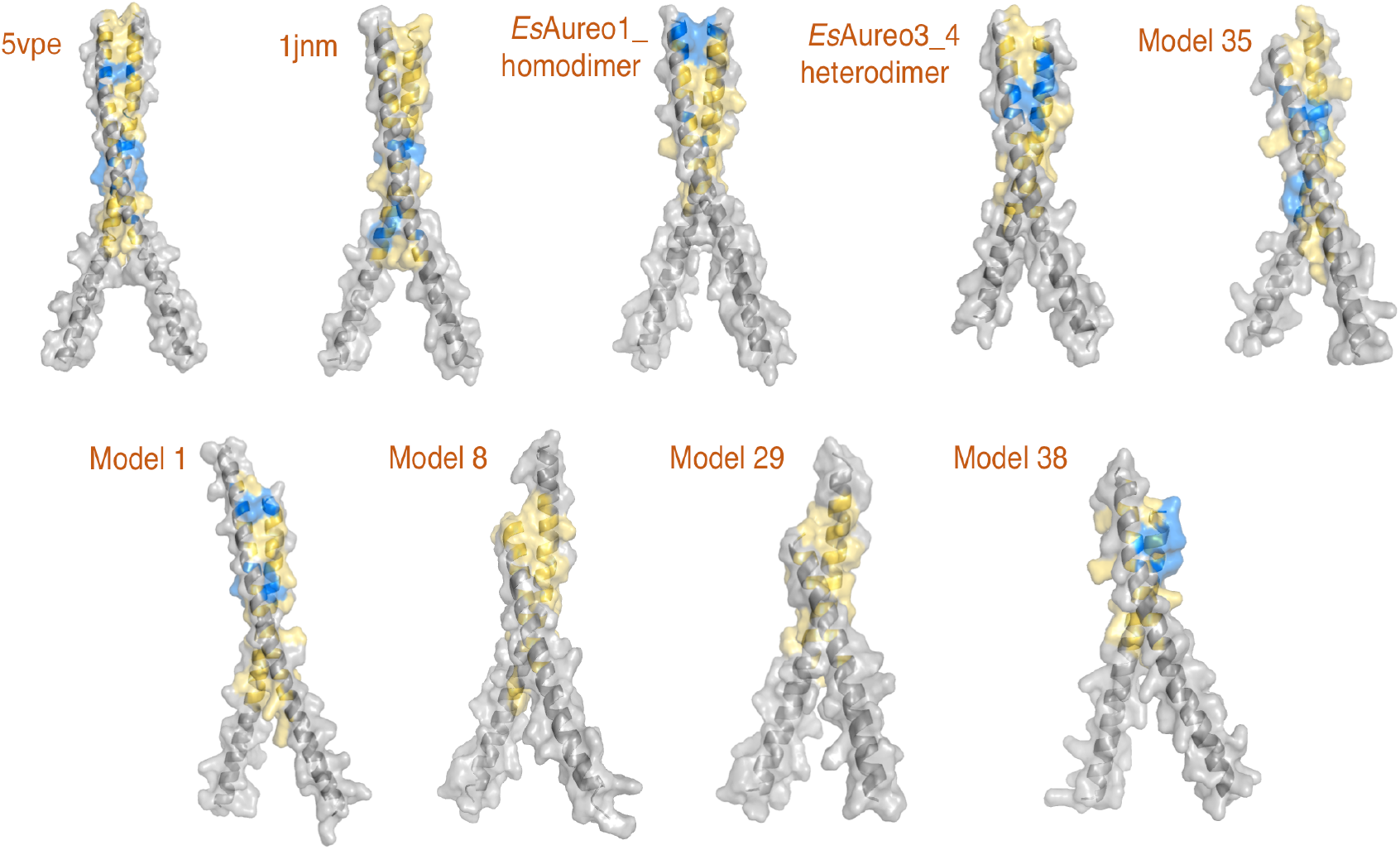
Interface analysis of selected dimers - 5vpe, 1jnm, EsAureo1_ homodimer, EsAureo3_4 heterodimer, Model_35, Model_1, Model_8, Model_29, and, Model_38. The interface residues are highlighted in yellow, whereas residues involved in hydrogen bonding / salt-bridge are highlighted in blue.

### Screening through heptad motifs

We next consider sequence composition of the heptads [**Figure 3**]. All bZIPs contain the required Asn and Arg in the DNA binding basic region, highlighted in red. It is well known that ‘g’ in second heptad and ‘e’ in the adjacent heptad is crucial for dimer compatibility. While electrostatic interactions strongly favor dimerization, repulsion will forbid the association of two monomeric units. In **Figure 3**, we delineate attraction in green, repulsion in red and non-electrostatic interaction (polar/non-polar) in black. As we can see, homodimerization of *Es*Aureo1, 2 and 3 are highly probable in this respect, while we see polar interactions in *Es*Aureo4. Alike *Es*Aureo4, we see polar interactions in the homodimeric structure of jun (PDB: 1jnm), as well. *Es*Aureo1, 3 and 4 are also predicted to be very stable according to KL divergence [**Table 2**]. *Es*Aureo3_4 heterodimer, which is predicted to be very stable according to ΔE_Sp_, also shows electrostatic attraction as well as polar interaction. This is comparable to the heterodimeric bZIP structure, 5vpe, comprising of fosB-junD. Very interestingly, in model_3, model_13 and model_23 there is at least one electrostatic repulsive force and should be unstable, as correctly pointed out by all measures. Further, in other models (model_9, model_19 and model_29) predicted to be unstable by measures from network and information theory, we can see unfavorable contact between charged and non-polar residues. Among others, model_20 and model_30, predicted to be stable by KL divergence and ΔE_Sp_, are indeed stabilized by electrostatic attractive forces. Overall, our results from heptad sequence analysis corroborates extremely well with the mathematical predictions.

**Figure 3:**
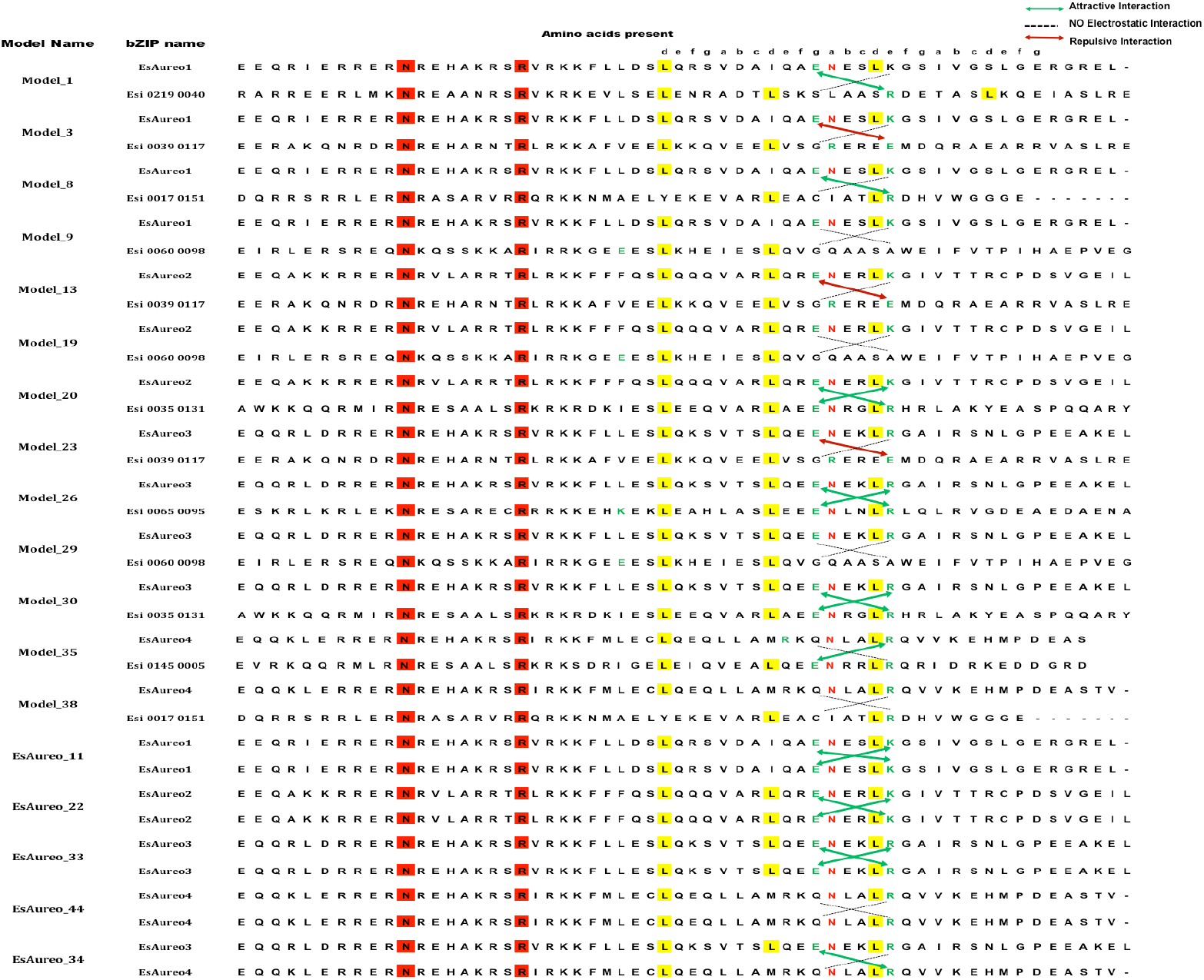
Screening of heptad sequences in selected model dimers. The electrostatic attractions and repulsions between ‘g’ and ‘e’ residues are marked with green and red respectively. The non-electrostatic forces including non-polar interactions are marked in black.

### Dynamics of the dimer interface

It has been claimed in earlier literature that a reasonable measure of information obtained from MD simulations to understand dimer stability can also be obtained by studying the average participation coefficient, P_av_, and specific difference of energy, ΔE_SP_ (Di Paola, Mei, et al., 2015)(Di Paola, Platania, et al., 2015). We conduct signature MD simulations on two EsAureo3_4 and model_29, which are predicted to be stable and unstable respectively in our investigations thus far. Following solvation, energy minimization and NVT/NPT equilibration, we conduct a production run of 20ns using GROMACS. The simulation trajectories are analyzed using the command line tools of GROMACS. **Figure 4** shows the simulation results comprising RMSD of C_α_ backbone from the initial structure as well as the inter-chain hydrogen bond interaction. In the 20ns simulation conducted at 300K, the RMSD values remain relatively high during the initial 4ns, for both cases. The RMSD for EsAureo3_4 later becomes relatively stable between 0.2nm and 0.4nm. In model_29, the RMSD varies between 0.4nm and 0.6nm. We, however, do not find much difference in inter-chain hydrogen bonding profile between EsAureo3_4 and model_29. Therefore, the overall RMSD trends over a timescale of 20ns shows beier stability of EsAureo3_4 over model_29, in agreement with all our other investigations. Obviously, these signature MD simulations are not a central focus of this paper, but have been conducted to complement the other exhaustive investigations undertaken herein.

**Figure 4:**
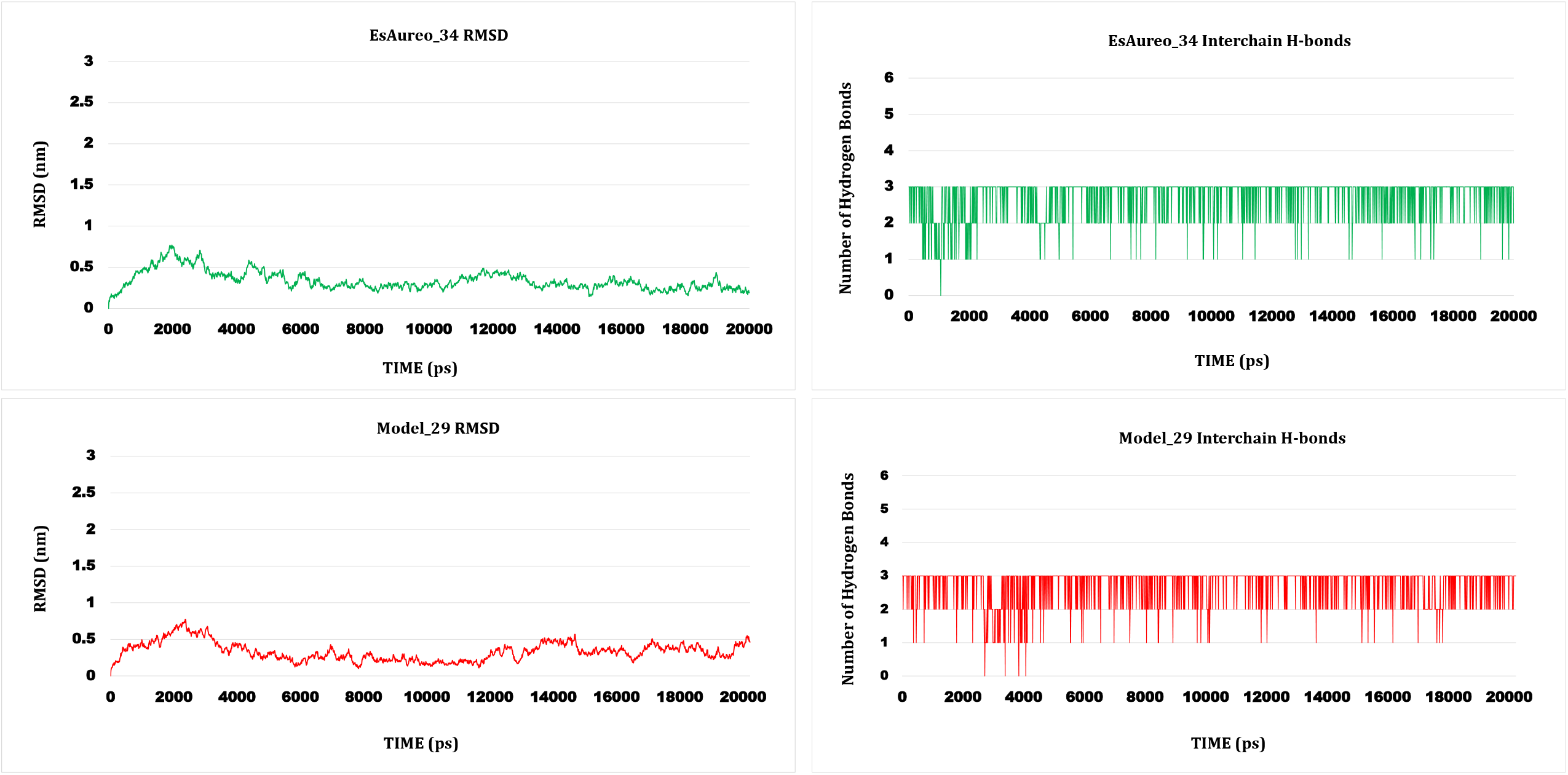
Signature MD simulations for EsAureo_34 and Model_29, showing the variations in RMSD as well as interchain hydrogen bonds.

## Discussion/Conclusion

The TF interaction network carries out the essential process of gene expression to determine cellular identity and function. With the availability of advanced computational methods in addition to high accuracy, low complexity and easy accumulation of ChIP-seq data, it is now possible to screen the combinatorial associations of TFs within a genome (Ye et al., 2017). One of the most interesting properties of TFs is their ability to share functional redundancy. In case of a loss-of-function scenario, TFs sharing a common phylogenetic clade with the missing TF can act interchangeably, indicating the robustness of the TF interaction network. Pharmacological modulation of a TF is daunting, considering the risks involved with the expression of hundreds of genes connected to the particular TF node. However, it has been shown in yeast that genetic disruption of a TF affects only 3% of the putative target (Wu & Li, 2015). Therefore, appraising paralog connectivity not only bears biochemical significance but is also profoundly important in pharmacology (Francois et al., 2018). In this paper, we focus on sensory TFs, aureochromes, which have the potential to regulate multiple cellular processes in response to blue light. Aureochromes are exclusively present in photosynthetic marine stramenopiles, where major LOV photoreceptors, phototropins are absent. We select *Ectocarpus siliculosus* as they harbor five potential aureochrome paralogs. The sensory nature of aureochromes calls for rapid transmission of signal to mediate gene activation/silencing — significant to understand natural phenomena and facilitating synthetic studies including optogenetics. The signal generated upon blue light absorption by C-terminal LOV sensor is allosterically transmiied to the bZIP domain (Banerjee et al., 2016) (Nakajima et al., 2021), located at the N-terminus. The 30-40 residues at the C-terminal of any bZIP protein generally constitute the zipper portion, which actually dimerizes with the corresponding zipper portion of another bZIP (Bader & Vogt, 2006a). Dimerization between bZIP TFs is a prerequisite to substrate DNA binding. Homo/hetero-dimerization between compatible bZIP partners is essential for choosing the substrate DNA sequence from a wide variety of substrate repertoire and thus to regulate diverse biological activities. In *Phaeodactylum tricornutum* (*Pt*), homo/heterodimerization among aureochrome bZIPs that eventually bind aureo-box sequence (TGACGT) is known experimentally. For example, *Pt*_aureochrome paralogs, *Pt*Aureo1a and 1c can undergo heterodimerization and display synergistic functions (Banerjee et al., 2016). Extended structural bioinformatic analyses further reveal the capability of *Pt*Aureo1a to heterodimerize with other paralogs, *Pt*Aureo1b, 1c as well as *Pt*Aureo2 — among which the last interaction is unstable in nature. Further, higher order structures are evident in the DNA binding assay *in vitro*, and might be associated with transcriptional regulation. While homo/hetero-dimerization is essential for the functioning of the bZIPs, concrete structural information behind higher order structure formation is awaited. In our all-theory study, we enquire whether aureochrome bZIP interactions are limited to aureochrome family or whether aureochrome bZIPs are compatible with other bZIP partners as well, within a given genome. We therefore collect all bZIP sequences (including those from aureochrome paralogs) from the fully-sequenced *Ectocarpus* genome, model all possible dimer complexes and analyze the data. Dimerization is interaction-specific and is largely dependent on amino acid sequence compositions. Again, certain dimers can prove to be energetically more stable than other dimers (Bader & Vogt, 2006a). This is the case for aureochrome bZIPs too. The availability of potential partners as well as the concentration of compatible partners in different sub cellular conditions determine the partnering specificity of bZIP dimers *in-vivo*. The essential question is - can aureochrome form homodimers as well as heterodimers with equal efficiency? Does the TF interaction network involving aureochromes include other bZIPs as wellti This study addresses this particular aspect using methods from structural bioinformatics, complex networks and information theory.

Interface analysis in terms of interface area, the gain in solvation energy, and, the number of hydrogen bonding/salt bridge interactions identify models with varied compatibility. The calculated interface area in the models vary from about 400Å^2^ to 900 Å^2^, indicating the difference in compatibility and stability of dimers. The results from analyses of interfaces is well supported by network-theoretic investigations on specific difference of energy and participation co-efficient, as well as information-theoretic measures like KL divergence and information centrality. As controls, we considered homo/ heterodimeric bZIP structures from PDB. All our measures predict compatibility and stability of the structures collected from PDB. With few exceptions, most of the *Ectocarpus* bZIPs demonstrate compatibility with the aureochrome bZIPs. The findings from all methods considered in this study are largely in good agreement with each other. The consensus obtained from our wide-ranging investigations allows us to select dimer models with higher and lower stability. Heptad composition and signature 20ns MD simulation runs further support the results of these investigations. Such ranking of aureochrome-based dimers in terms of compatibility and stability would surely facilitate the seamless design of future experiments.

## Statements

### Statement of ethics

This is a purely computational study. No experiments on animal or human subjects are involved and therefore ethics approval should not be required.

### Author Contribution

Madhurima Khamaru, Deep Nath and Devrani Mitra were involved with the generation and analysis of data. Madhurima Khamaru worked on modeling, interface analysis, and MD simulations while Deep Nath worked on complex networks and information theory. Devrani Mitra and Soumen Roy designed and supervised the research. Devrani Mitra and Soumen Roy wrote the manuscript with the help of Madhurima Khamaru and Deep Nath.

### Data availability statement

This is a purely theoretical paper and has no experimental data. All data generated or analyzed during this study are included in the main text article as well as in supplementary information. Further enquiries can be directed to the corresponding authors.

