## Supplementary material for "Assessing combinatorial diversity of aureochrome bZIPs through genome-wide screening": Supplmentary Information

This file contains **Table SI_1 and Table SI_2**

**Table SI_1 is constituted from the following tables:**

Table-1. a) Interface Analysis Data of all the Modelled Heterodimers of AureobZIPs and bZIPs from *Ectocarpus siliculosus*

Table-1. b) Interface Analysis Data of all the Modelled Heterodimers of AureobZIPs of *Ectocarpus siliculosus*

Table-1. c) Interface Analysis Data of all the Modelled Homodimers of AureobZIPs from *Ectocarpus siliculosus* and the interface analysis data of the crystal structures of known bZIP homo and heterodimers

**Table SI_2:** Relevant data to calculate the specific difference of energy (ΔESP), average participation coefficient (Pav), Kullback-Leibler divergence (KL), and, relative information centrality (Ir) for all possible dimers.

| Protein A | Protein B | Name of dimer | Interface Area ($A^2$) | Delta G | H-bond | salt bridge |
|---|---|---|---|---|---|---|
| | Esi_0219_0040 | Model1 | 871 | -17.2 | 1 A:GLU 39[ OE1] 3.45 B:ARG 45[ NH1]<br>2 A:ASN 40[ OD1] 3.82 B:SER 40[ OG ]<br>3 A:ASN 40[ OD1] 3.76 B:SER 44[ OG ]<br>4 A:GLU 53[ OE1] 3.15 B:ARG 59[ NH1] | 1 A:ARG 54[ NE ] 3.58 B:GLU 54[ OE1]<br>2 A:GLU 39[ OE1] 3.45 B:ARG 45[ NH1]<br>3 A:GLU 53[ OE1] 3.15 B:ARG 59[ NH1] |
| | Esi_0113_0081 | Model2 | 876.4 | -16.3 | 1 A:ASN 40[ ND2] 2.88 B:ASN 41[ OD1]<br>2 A:GLN 30[ OE1] 2.75 B:ARG 26[ NH2]<br>3 A:ILE 36[ O ] 3.10 B:ASN 41[ ND2]<br>4 A:SER 46[ OG ] 2.92 B:ASN 48[ ND2] | NA |
| | Esi_0039_0117 | Model3 | 515 | -11 | 1 A:GLU 39[ OE1] 2.75 B:ARG 41[ NH2] | 1 A:GLU 39[ OE1] 2.75 B:ARG 41[ NH2] |
| | Esi_0199_0057 | Model4 | 727.1 | -12.6 | 1 A:ARG 54[ NH1] 3.77 B:THR 53[ O ]<br>2 A:GLU 39[ OE1] 2.79 B:ARG 45[ NH2] | 1 A:ARG 54[ NH1] 3.77 B:THR 53[ O ]<br>2 A:GLU 39[ OE1] 2.79 B:ARG 45[ NH2] |
| EsAureo1 | Esi_0145_0005 | Model5 | 853.5 | -16.1 | 1 A:ASN 40[ ND2] 2.79 B:ASN 41[ OD1]<br>2 A:GLN 30[ OE1] 2.76 B:ARG 26[ NH2]<br>3 A:ILE 36[ O ] 3.11 B:ASN 41[ ND2] | 1 A:ARG 54[ NE ] 3.98 B:ASP 54[ OD2]<br>2 A:ARG 54[ NH1] 3.86 B:ASP 54[ OD2]<br>3 A:ARG 54[ NH2] 3.18 B:ASP 54[ OD2] |
| | Esi_0065_0095 | Model6 | 787.5 | -14.2 | 1 A:ASN 40[ ND2] 3.80 B:GLU 40[ OE1]<br>2 A:ASN 40[ ND2] 2.86 B:ASN 41[ OD1]<br>3 A:ILE 36[ O ] 3.15 B:ASN 41[ ND2]<br>4 A:GLU 39[ OE1] 3.02 B:ARG 45[ NH1] | 1 A:GLU 39[ OE1] 3.02 B:ARG 45[ NH1] |
| | Esi_0211_0027 | Model7 | 696.70 | -13.2 | 1 A:ALA 50[ O ] 2.46 B:ARG 54[HH12] | 1 A:ALA 50[ O ] 3.44 B:ARG 54[ NH1] |
| | Esi_0017_0151 | Model8 | 549.70 | -11.7 | NA | NA |
| | Esi_0060_0098 | Model9 | 447.40 | -10.7 | NA | NA |
| | Esi_0035_0131 | Model10 | 856.9 | -17.2 | 1 A:ASN 40[ ND2] 3.76 B:GLU 40[ OE1]<br>2 A:ASN 40[ ND2] 2.77 B:ASN 41[ OD1]<br>3 A:ARG 54[ NE ] 2.97 B:SER 54[ OG ]<br>4 A:ILE 36[ O ] 3.08 B:ASN 41[ ND2]<br>5 A:GLU 39[ OE1] 3.66 B:ASN 41[ ND2] | NA |
| | Esi_0219_0040 | Model11 | 577.10 | -11.2 | 1 A:GLU 39[ OE2] 1.74 D:ARG 44[HH12] | 1 A:GLU 39[ OE2] 2.64 D:ARG 44[ NH1] |
| | Esi_0113_0081 | Model12 | 723.2 | -11.2 | 1 A:ASN 40[ ND2] 2.86 B:ASN 41[ OD1]<br>2 A:GLN 30[ OE1] 2.75 B:ARG 26[ NH2]<br>3 A:LEU 36[ O ] 3.06 B:ASN 41[ ND2]<br>4 A:LEU 43[ O ] 3.08 B:ASN 48[ ND2] | NA |
| | Esi_0039_0117 | Model13 | 490.1 | -11.5 | NA | NA |
| | Esi_0199_0057 | Model14 | 664.6 | -15 | 1 A:ASN 40[ ND2] 3.83 B:LEU 41[ O ] | NA |
| | Esi_0145_0005 | Model15 | 750.1 | -14.8 | 1 A:ASN 40[ ND2] 2.80 B:ASN 41[ OD1]<br>2 A:GLN 30[ OE1] 2.74 B:ARG 26[ NH2]<br>3 A:LEU 36[ O ] 3.05 B:ASN 41[ ND2] | NA |
| EsAureo2 | Esi_0065_0095 | Model16 | 736.3 | -12.9 | 1 B:GLN 32[ NE2] 3.14 A:GLU 38[ OE2]<br>2 B:ASN 40[ ND2] 3.04 A:LEU 37[ O ]<br>3 B:GLU 39[ OE2] 3.23 A:ARG 45[ NH1]<br>4 B:ASN 40[ OD1] 2.83 A:ASN 41[ ND2] | 1 B:GLU 39[ OE2] 3.23 A:ARG 45[ NH1] |
| | Esi_0211_0027 | Model17 | 646.5 | -15.1 | 1 A:VAL 33[ N ] 3.66 B:SER 34[ OG ] | NA |
| | Esi_0017_0151 | Model18 | 591.8 | -10.1 | 1 A:TYR 30[ HH ] 2.05 B:LEU 29[ O ]<br>2 A:GLU 38[ OE1] 1.95 B:ARG 35[HH12]<br>3 A:GLU 38[ OE2] 2.22 B:ARG 35[HH11] | 1 A:GLU 38[ OE1] 2.86 B:ARG 35[ NH1]<br>2 A:GLU 38[ OE2] 2.85 B:ARG 35[ NH1] |
| | Esi_0060_0098 | Model19 | 461.4 | -6.8 | NA | NA |
| | Esi_0035_0131 | Model20 | 674.5 | -14.3 | 1 A:ASN 40[HD21] 2.50 D:GLU 40[ OE1]<br>2 A:ASN 40[HD22] 2.18 D:LEU 37[ O ] | NA |
| | Esi_0219_0040 | Model21 | 638.6 | -9.3 | 1 A:GLU 39[ OE2] 1.63 D:ARG 44[HH11] | 1 A:GLU 39[ OE2] 3.92 D:ARG 44[ NE ]<br>2 A:GLU 39[ OE2] 2.61 D:ARG 44[ NH1] |
| | Esi_0113_0081 | Model22 | 817.3 | -12.4 | 1 A:ASN 40[ ND2] 2.72 B:ASN 41[ OD1]<br>2 A:ARG 44[ NH1] 2.53 B:GLU 40[ OE1]<br>3 A:ASN 50[ ND2] 2.90 B:ASN 48[ OD1]<br>4 A:GLN 30[ OE1] 2.72 B:ARG 26[ NH2]<br>5 A:LEU 36[ O ] 3.01 B:ASN 41[ ND2] | 1 A:ARG 44[ NH1] 2.53 B:GLU 40[ OE1]<br>2 A:ARG 44[ NH2] 3.97 B:GLU 40[ OE1] |
| EsAureo3 | Esi_0039_0117 | Model23 | 521.8 | -11 | 1 A:GLU 39[ OE1] 2.78 B:ARG 41[ NH2] | 1 A:GLU 39[ OE1] 2.78 B:ARG 41[ NH2] |
| | Esi_0199_0057 | Model24 | 705 | -14.6 | 1 A:ARG 44[ NH1] 3.14 B:GLU 44[ OE2]<br>2 A:ASN 50[ ND2] 3.78 B:SER 52[ O ] | 1 A:ARG 44[ NE ] 3.91 B:GLU 44[ OE2]<br>2 A:ARG 44[ NH1] 3.14 B:GLU 44[ OE2] |
| | Esi_0145_0005 | Model25 | 803.1 | -14 | 1 A:ASN 40[ ND2] 2.79 B:ASN 41[ OD1]<br>2 A:ARG 44[ NH1] 3.20 B:GLU 40[ OE2]<br>3 A:GLN 30[ OE1] 2.72 B:ARG 26[ NH2]<br>4 A:LEU 36[ O ] 3.01 B:ASN 41[ ND2] | 1 A:ARG 44[ NE ] 3.95 B:GLU 40[ OE2]<br>2 A:ARG 44[ NH1] 3.20 B:GLU 40[ OE2] |
| | Esi_0065_0095 | Model26 | 801.2 | -13 | 1 A:ASN 40[ ND2] 3.62 B:GLU 40[ OE1]<br>2 A:ASN 40[ ND2] 2.82 B:ASN 41[ OD1] | 1 A:ARG 44[ NH1] 3.51 B:GLU 40[ OE1]<br>2 A:GLU 39[ OE1] 2.82 B:ARG 45[ NH1] |

| | | | | | | |
|---|---|---|---|---|---|---|
| | | | | | 3 A:ARG 44[ NH1] 3.51 B:GLU 40[ OE1]<br>4 A:LEU 36[ O ] 3.05 B:ASN 41[ ND2]<br>5 A:GLU 39[ OE1] 2.82 B:ARG 45[ NH1] | |
| | Esi_0211_0027 | Model27 | 637.1 | -14.9 | 1 A:VAL 33[ N ] 3.65 B:SER 34[ OG ]<br>2 A:ASN 40[ OD1] 3.18 B:HIS 41[ ND1] | NA |
| | Esi_0017_0151 | Model28 | 643.8 | -13.2 | 1 A:GLU 39[ OE1] 2.97 B:ARG 45[ NH1] | 1 A:GLU 39[ OE1] 2.97 B:ARG 45[ NH1] |
| | Esi_0060_0098 | Model29 | 389.3 | 8.6 | NA | NA |
| | Esi_0035_0131 | Model30 | 809.5 | -14.9 | 1 A:ASN 40[ ND2] 3.66 B:GLU 40[ OE1]<br>2 A:ASN 40[ ND2] 2.72 B:ASN 41[ OD1]<br>3 A:ASN 50[ ND2] 3.05 B:TYR 51[ O ]<br>4 A:LEU 36[ O ] 3.06 B:ASN 41[ ND2]<br>5 A:GLU 39[ OE1] 3.86 B:ASN 41[ ND2] | 1 A:ARG 44[ NH1] 3.91 B:GLU 40[ OE1] |
| EsAureo4 | Esi_0219_0040 | Model31 | 657.9 | -8.4 | 1 C:GLN 39[ OE1] 2.27 D:ARG 44[ HE ] | NA |
| | Esi_0113_0081 | Model32 | 850.9 | -13 | 1 A:ARG 44[ NH1] 2.61 B:GLU 40[ OE2]<br>2 A:ASN 40[ ND2] 2.81 B:ASN 41[ OD1]<br>3 A:GLN 30[ OE1] 2.71 B:ARG 26[ NH2]<br>4 A:MET 36[ O ] 3.04 B:ASN 41[ ND2] | 1 A:ARG 44[ NH1] 2.61 B:GLU 40[ OE2] |
| | Esi_0039_0117 | Model33 | 570.2 | -9.1 | 1 A:ARG 40[HH12] 1.69 D:ALA 35[ O ]<br>2 A:ARG 40[HH22] 1.75 D:ALA 35[ O ]<br>3 A:ARG 40[ HE ] 1.93 D:MET 36[ O ]<br>4 A:ARG 40[HH21] 2.30 D:MET 36[ SD ]<br>5 A:GLY 39[ O ] 2.46 D:ASN 40[HD21] | NA |
| | Esi_0199_0057 | Model34 | 643.5 | -14.1 | 1 B:GLU 44[ OE2] 1.69 C:ARG 44[HH12] | 1 B:GLU 44[ OE2] 2.72 C:ARG 44[ NH1]<br>2 B:GLU 44[ OE2] 3.50 C:ARG 44[ NE ] |
| | Esi_0145_0005 | Model35 | 854.6 | -15.5 | 1 D:ARG 44[ NH1] 3.06 E:GLU 40[ OE2]<br>2 D:ASN 40[ ND2] 2.83 E:ASN 41[ OD1]<br>3 D:GLN 30[ OE1] 2.75 E:ARG 26[ NH2]<br>4 D:MET 36[ O ] 3.26 E:ASN 41[ ND2]<br>5 D:GLN 39[ OE1] 2.85 E:ASN 41[ ND2] | 1 D:ARG 44[ NH1] 3.06 E:GLU 40[ OE2] |
| | Esi_0065_0095 | Model36 | 779.1 | -15.5 | 1 B:ASN 40[ ND2] 3.06 A:LEU 37[ O ]<br>2 B:ARG 44[ NH1] 3.03 A:GLU 40[ OE1]<br>3 B:ASN 40[ OD1] 2.79 A:ASN 41[ ND2] | 1 B:ARG 44[ NH1] 3.03 A:GLU 40[ OE1] |
| | Esi_0211_0027 | Model37 | 742.3 | -18.3 | 1 E:ASN 40[ ND2] 3.39 D:LEU 38[ O ]<br>2 E:ARG 44[ N ] 3.86 D:MET 45[ SD ] | NA |
| | Esi_0017_0151 | Model38 | 511.2 | -12.3 | 1 E:MET 36[ SD ] 3.41 F:GLU 38[ N ]<br>2 E:GLN 39[ OE1] 3.27 F:ARG 45[ NH1] | NA |
| | Esi_0060_0098 | Model39 | 611.2 | -10 | 1 E:ARG 37[ NH2] 2.92 D:GLU 33[ OE2]<br>2 E:ASN 40[ ND2] 3.28 D:LEU 37[ O ]<br>3 E:GLN 39[ OE1] 3.02 D:GLN 41[ NE2] | 1 E:ARG 37[ NH2] 2.92 D:GLU 33[ OE2] |
| | Esi_0035_0131 | Model40 | 821.2 | -15.2 | 1 D:ASN 40[ ND2] 3.75 E:GLU 40[ OE1]<br>2 D:ARG 44[ NH1] 3.74 E:GLU 40[ OE1]<br>3 D:ASN 40[ ND2] 2.75 E:ASN 41[ OD1]<br>4 D:MET 36[ O ] 3.21 E:ASN 41[ ND2]<br>5 D:GLN 39[ OE1] 2.86 E:ASN 41[ ND2] | 1 D:ARG 44[ NH1] 3.74 E:GLU 40[ OE1] |

### Table-1. a) Interface Analysis Data of all the Modelled Heterodimers of AureobZIPs and bZIPs from *Ectocarpus siliculosus*

| Protein A | Protein B | Name of dimer | Interface Area ($A^2$) | Delta G | H-bond | salt bridge |
|---|---|---|---|---|---|---|
| EsAureo1 | EsAureo2 | EsAureo12 | 747.1 | -14.4 | 1 A:ASN 40[ ND2] 2.84 B:ASN 40[ OD1]<br>2 A:ARG 54[ NH1] 3.83 B:ILE 46[ O ]<br>3 A:ILE 36[ O ] 3.13 B:ASN 40[ ND2] | 1 A:ARG 54[ NH1] 3.95 B:VAL 47[ OXT] |
| | EsAureo3 | EsAureo13 | 697.7 | -15.1 | 1 A:ASN 40[ ND2] 3.80 B:GLU 39[ OE1]<br>2 A:ASN 40[ ND2] 2.79 B:ASN 40[ OD1]<br>3 A:ILE 36[ O ] 3.12 B:ASN 40[ ND2] | NA |
| | EsAureo4 | EsAureo14 | 786 | -13.7 | 1 B:ASN 40[ ND2] 3.12 A:ILE 36[ O ]<br>2 B:ARG 44[ NH1] 2.93 A:GLU 39[ OE1]<br>3 B:ASN 40[ OD1] 2.84 A:ASN 40[ ND2]<br>4 B:VAL 46[ O ] 3.89 A:ARG 54[ NH1]<br>5 B:LYS 48[ O ] 3.77 A:ARG 54[ NH1] | 1 B:ARG 44[ NH1] 2.93 A:GLU 39[ OE1]<br>2 B:LYS 48[ O ] 3.77 A:ARG 54[ NH1] |
| EsAureo2 | EsAureo1 | EsAureo12 | 747.1 | -14.4 | 1 A:ASN 40[ ND2] 2.84 B:ASN 40[ OD1] | 1 A:ARG 54[ NH1] 3.95 B:VAL 47[ OXT] |

| | | | | | | |
|---|---|---|---|---|---|---|
| | | | | | 2 A:ARG 54[ NH1] 3.83 B:ILE 46[ O ]<br>3 A:ILE 36[ O ] 3.13 B:ASN 40[ ND2] | |
| | **EsAureo3** | **EsAureo23** | **742** | **-14.7** | 1 B:ASN 40[ ND2] 3.04 A:LEU 36[ O ]<br>2 B:GLU 39[ OE2] 3.20 A:ARG 44[ NH1]<br>3 B:ASN 40[ OD1] 2.81 A:ASN 40[ ND2] | 1 B:GLU 39[ OE2] 3.97 A:ARG 44[ NE ]<br>2 B:GLU 39[ OE2] 3.20 A:ARG 44[ NH1] |
| | **EsAureo4** | **EsAureo24** | **623.3** | **-10.3** | 1 A:ARG 37[HH12] 1.87 B:GLN 32[ OE1]<br>2 A:ARG 37[HH22] 2.04 B:GLN 32[ OE1]<br>3 A:ASN 40[HD22] 2.36 B:LEU 36[ O ]<br>4 A:ASN 40[ OD1] 1.88 B:ASN 40[HD21] | **NA** |
| **EsAureo3** | **EsAureo1** | **EsAureo13** | **697.7** | **-15.1** | 1 A:ASN 40[ ND2] 3.80 B:GLU 39[ OE1]<br>2 A:ASN 40[ ND2] 2.79 B:ASN 40[ OD1]<br>3 A:ILE 36[ O ] 3.12 B:ASN 40[ ND2] | **NA** |
| | **EsAureo2** | **EsAureo23** | **742** | **-14.7** | 1 B:ASN 40[ ND2] 3.04 A:LEU 36[ O ]<br>2 B:GLU 39[ OE2] 3.20 A:ARG 44[ NH1]<br>3 B:ASN 40[ OD1] 2.81 A:ASN 40[ ND2] | 1 B:GLU 39[ OE2] 3.97 A:ARG 44[ NE ]<br>2 B:GLU 39[ OE2] 3.20 A:ARG 44[ NH1] |
| | **EsAureo4** | **EsAureo34** | **765.3** | **-16** | 1 A:GLN 37[ N ] 3.61 B:MET 36[ SD ]<br>2 A:ASN 40[ ND2] 2.76 B:ASN 40[ OD1]<br>3 A:LEU 36[ O ] 3.04 B:ASN 40[ ND2]<br>4 A:GLU 39[ OE1] 2.89 B:ARG 44[ NH1] | 1 A:GLU 39[ OE1] 2.89 B:ARG 44[ NH1] |
| **EsAureo4** | **EsAureo1** | **EsAureo14** | **786** | **-13.7** | 1 B:ASN 40[ ND2] 3.12 A:ILE 36[ O ]<br>2 B:ARG 44[ NH1] 2.93 A:GLU 39[ OE1]<br>3 B:ASN 40[ OD1] 2.84 A:ASN 40[ ND2]<br>4 B:VAL 46[ O ] 3.89 A:ARG 54[ NH1]<br>5 B:LYS 48[ O ] 3.77 A:ARG 54[ NH1] | 1 B:ARG 44[ NH1] 2.93 A:GLU 39[ OE1]<br>2 B:LYS 48[ O ] 3.77 A:ARG 54[ NH1] |
| | **EsAureo2** | **EsAureo24** | **798.1** | **-15** | 1 A:ARG 37[ NH1] 2.86 B:GLN 32[ OE1]<br>2 A:ASN 40[ ND2] 3.34 B:LEU 36[ O ]<br>3 A:ASN 40[ ND2] 3.42 B:GLU 39[ OE2]<br>4 A:ASN 40[ OD1] 2.84 B:ASN 40[ ND2] | **NA** |
| | **EsAureo3** | **EsAureo34** | **765.3** | **-16** | 1 A:GLN 37[ N ] 3.61 B:MET 36[ SD ]<br>2 A:ASN 40[ ND2] 2.76 B:ASN 40[ OD1]<br>3 A:LEU 36[ O ] 3.04 B:ASN 40[ ND2]<br>4 A:GLU 39[ OE1] 2.89 B:ARG 44[ NH1] | 1 A:GLU 39[ OE1] 2.89 B:ARG 44[ NH1] |

**Table-1. b) Interface Analysis Data of all the Modelled Heterodimers of AureobZIPs of *Ectocarpus siliculosus***

| **Name of dimer** | **Interface Area ($A^2$)** | **Delta G** | **H-bond** | **salt bridge** |
|---|---|---|---|---|
| **EsAu1_homodimer** | **790.5** | **-10.3** | 1 A:ASN 40[HD21] 2.26 B:ILE 36[ O ]<br>2 A:SER 50[ OG ] 2.71 B:SER 50[ OG ]<br>3 A:ARG 54[ HE ] 2.25 B:SER 50[ O ] | 1 A:ARG 54[ NE ] 2.69 B:GLU 53[ OE1]<br>2 A:ARG 54[ NH1] 3.15 B:GLU 53[ OE1]<br>3 A:ARG 54[ NH2] 3.70 B:GLU 53[ OE2]<br>4 A:GLU 53[ OE1] 3.15 B:ARG 54[ NH2] |
| **EsAu2_homodimer** | **660** | **-13.7** | 1 A:ASN 40[HD21] 2.00 B:LEU 36[ O ]<br>2 A:ASN 40[ OD1] 1.80 B:ASN 40[HD22] | 1 A:LYS 44[ NZ ] 3.38 B:GLU 39[ OE2] |
| **EsAu3_homodimer** | **703.3** | **-13.7** | 1 A:ASN 40[HD21] 2.11 B:LEU 36[ O ]<br>2 A:ASN 40[ OD1] 2.02 B:ASN 40[HD21] | **NA** |
| **EsAu4_homodimer** | **724.3** | **-15.6** | 1 C:GLN 39[ OE1] 2.17 D:ASN 40[HD21] | **NA** |
| **5VPE** | **1181.8** | **-21.7** | 1 B:ARG 292[HH21] 2.40 A:GLN 184[ OE1]<br>2 B:LYS 299[ HZ1] 2.43 A:GLU 191[ OE2]<br>3 B:ARG 318[HH11] 1.71 A:GLU 200[ OE2]<br>4 B:GLN 306[ OE1] 2.00 A:LYS 194[ HZ1]<br>5 B:GLN 320[ OE1] 1.98 A:LYS 208[ HZ3] | 1 B:LYS 299[ NZ ] 2.76 A:GLU 191[ OE2]<br>2 B:LYS 304[ NZ ] 3.33 A:GLU 186[ OE2]<br>3 B:ARG 318[ NH1] 2.27 A:GLU 200[ OE2]<br>4 B:ARG 318[ NH1] 3.46 A:GLU 200[ OE1] |
| **1JNM** | **1000.6** | **-14.9** | 1 B:ARG 276[ NH1] 3.12 A:GLU 281[ OE2]<br>2 B:ASN 291[ ND2] 2.84 A:GLN 290[ OE1]<br>3 B:GLN 290[ NE2] 3.23 A:ASN 291[ OD1] | 1 B:ARG 276[ NH1] 3.12 A:GLU 281[ OE2]<br>2 B:GLU 281[ OE1] 2.36 A:ARG 276[ NH2] |

| | | | | |
|---|---|---|---|---|
| | | | 4 B:GLU 281[ OE1] 2.36 A:ARG 276[ NH2]<br>5 B:LEU 287[ O ] 3.69 A:ASN 291[ ND2]<br>6 B:ASN 291[ OD1] 2.35 A:ASN 291[ ND2] | |
| **1fos** | **1188.5** | **-18.7** | 1 F:ARG 285[ NH2] 2.99 E:GLN 166[ OE1]<br>2 F:LYS 292[ NZ ] 3.02 E:GLU 173[ OE2]<br>3 F:LYS 297[ NZ ] 3.11 E:GLU 168[ OE2]<br>4 F:ARG 311[ NH2] 3.71 E:GLU 182[ OE1]<br>5 F:LYS 318[ NZ ] 2.90 E:GLU 189[ OE1]<br>6 F:GLN 299[ OE1] 2.94 E:LYS 176[ NZ ]<br>7 F:ASN 300[ OD1] 2.99 E:LYS 176[ N ]<br>8 F:GLN 313[ OE1] 3.24 E:LYS 190[ NZ ] | 1 F:LYS 292[ NZ ] 3.02 E:GLU 173[ OE2]<br>2 F:LYS 297[ NZ ] 3.11 E:GLU 168[ OE2]<br>3 F:ARG 311[ NE ] 3.37 E:GLU 182[ OE2]<br>4 F:ARG 311[ NH1] 3.34 E:GLU 182[ OE2]<br>5 F:ARG 311[ NH2] 3.71 E:GLU 182[ OE1]<br>6 F:ARG 311[ NH2] 3.62 E:GLU 182[ OE2]<br>7 F:LYS 318[ NZ ] 2.90 E:GLU 189[ OE1] |
| **JUN** | **923.5** | **-21** | 1 A:CYS 273[ SG ] 2.02 B:CYS 273[ SG ] | 1 A:CYS 273[ SG ] 2.02 B:CYS 273[ SG ] |

**Table-1. c) Interface Analysis Data of all the Modelled Homodimers of AureobZIPs from *Ectocarpus siliculosus* and the interface analysis data of the crystal structures of known bZIP homo and heterodimers**

*** Note: The residue numbers of the modeled structures differ from those in original sequences. This is due to modeling against the selected portions of bZIPs.**

| #model | $N_D$ | $I_D$ | $I_{M1}$ | $I_{M2}$ | $I_r$ | $P_{av}$ | $E_D$ | $E_{M1}$ | $E_{M2}$ | $\Delta E_{SP}$ | KL div. |
|---|---|---|---|---|---|---|---|---|---|---|---|
| 5vpe | 134 | 0.01541 | 0.02453 | 0.02499 | 0.68881 | 0.14243 | 291.93013 | 136.20560 | 136.17036 | 0.14593 | 2.07409 |
| 1jnm | 113 | 0.01483 | 0.02448 | 0.02284 | 0.68660 | 0.14620 | 241.33107 | 109.87945 | 110.78964 | 0.18285 | 1.63791 |
| 1jun | 86 | 0.03173 | 0.05067 | 0.05067 | 0.68690 | 0.17506 | 187.29721 | 85.24337 | 85.24337 | 0.19547 | 2.82295 |
| 1fos | 117 | 0.01417 | 0.02174 | 0.02218 | 0.67737 | 0.15265 | 249.24628 | 115.02250 | 112.60333 | 0.18479 | 1.87273 |
| EsAu1_1 | 108 | 0.02004 | 0.03723 | 0.03657 | 0.72846 | 0.14307 | 234.46554 | 109.09168 | 109.51106 | 0.14688 | 2.20119 |
| EsAu2_2 | 94 | 0.01591 | 0.03292 | 0.03245 | 0.75662 | 0.11181 | 195.24344 | 91.58499 | 91.54180 | 0.12890 | 1.33022 |
| EsAu3_3 | 96 | 0.02263 | 0.04556 | 0.04451 | 0.74875 | 0.12371 | 206.16626 | 96.68042 | 97.32895 | 0.12663 | 1.88449 |
| EsAu4_4 | 92 | 0.02560 | 0.04674 | 0.04788 | 0.72944 | 0.12539 | 198.15428 | 92.37580 | 93.05535 | 0.13830 | 1.91047 |
| EsAu1_2 | 99 | 0.01631 | 0.02627 | 0.03552 | 0.73604 | 0.11805 | 206.87713 | 106.04113 | 87.62970 | 0.13340 | 1.39643 |
| EsAu1_3 | 101 | 0.01594 | 0.02627 | 0.03317 | 0.73183 | 0.10830 | 211.16088 | 106.04113 | 92.11548 | 0.12876 | 1.34515 |
| EsAu1_4 | 100 | 0.01642 | 0.02627 | 0.03394 | 0.72729 | 0.12651 | 210.39719 | 106.04113 | 89.53297 | 0.14823 | 1.45623 |
| EsAu2_3 | 95 | 0.01708 | 0.02943 | 0.03552 | 0.73703 | 0.12268 | 198.99185 | 97.79645 | 87.62970 | 0.14280 | 1.47127 |
| EsAu2_4 | 89 | 0.02491 | 0.05568 | 0.04661 | 0.75648 | 0.11269 | 190.05614 | 84.32018 | 94.83992 | 0.12243 | 1.67723 |
| EsAu3_4 | 96 | 0.01713 | 0.02943 | 0.03394 | 0.72968 | 0.12957 | 202.52961 | 97.79645 | 89.53297 | 0.15834 | 1.55752 |
| model_01 | 119 | 0.01236 | 0.02554 | 0.01805 | 0.71645 | 0.10523 | 247.90011 | 105.51705 | 126.79202 | 0.13102 | 1.21647 |
| model_02 | 107 | 0.01508 | 0.02627 | 0.02647 | 0.71407 | 0.13187 | 225.30154 | 106.04113 | 103.47419 | 0.14754 | 1.51889 |
| model_03 | 93 | 0.01613 | 0.02627 | 0.04422 | 0.77117 | 0.08704 | 190.79144 | 106.04113 | 75.17265 | 0.10299 | 1.01498 |
| model_04 | 102 | 0.01541 | 0.02627 | 0.03092 | 0.73055 | 0.12333 | 213.66401 | 106.04113 | 93.06699 | 0.14271 | 1.47358 |
| model_05 | 108 | 0.01459 | 0.02627 | 0.02576 | 0.71958 | 0.11852 | 227.03854 | 106.04113 | 105.72871 | 0.14138 | 1.40576 |
| model_06 | 103 | 0.01561 | 0.02627 | 0.03050 | 0.72503 | 0.11664 | 216.11554 | 106.04113 | 95.56694 | 0.14085 | 1.43219 |
| Model_07 | 102 | 0.02240 | 0.04337 | 0.03598 | 0.71771 | 0.12580 | 220.75539 | 96.83387 | 109.26466 | 0.14369 | 1.77896 |
| Model_08 | 99 | 0.01939 | 0.04890 | 0.03552 | 0.77032 | 0.09801 | 209.15321 | 90.31312 | 109.04983 | 0.09889 | 1.45705 |
| Model_09 | 90 | 0.02361 | 0.03831 | 0.07194 | 0.78585 | 0.07539 | 189.91835 | 110.22041 | 71.74608 | 0.08835 | 1.18139 |
| model_10 | 108 | 0.01478 | 0.02627 | 0.02562 | 0.71517 | 0.11972 | 226.81932 | 106.04113 | 105.44150 | 0.14201 | 1.44944 |
| Model_11 | 96 | 0.02215 | 0.04831 | 0.03917 | 0.74680 | 0.11620 | 206.38243 | 95.37433 | 98.27628 | 0.13262 | 1.71205 |
| model_12 | 101 | 0.01537 | 0.03143 | 0.02647 | 0.73454 | 0.10521 | 209.71506 | 93.59330 | 103.47419 | 0.12522 | 1.26912 |
| model_13 | 87 | 0.01730 | 0.03143 | 0.04422 | 0.77132 | 0.08290 | 177.70944 | 93.59330 | 75.17265 | 0.10280 | 1.00612 |
| model_14 | 96 | 0.01577 | 0.03143 | 0.03092 | 0.74707 | 0.10338 | 198.71341 | 93.59330 | 93.06699 | 0.12555 | 1.28670 |
| model_15 | 102 | 0.01486 | 0.03143 | 0.02576 | 0.74016 | 0.10974 | 212.34984 | 93.59330 | 105.72871 | 0.12772 | 1.32013 |
| model_16 | 95 | 0.01665 | 0.02879 | 0.03552 | 0.74110 | 0.11967 | 198.48560 | 96.98248 | 87.62970 | 0.14604 | 1.41375 |
| model_17 | 93 | 0.01715 | 0.03143 | 0.03573 | 0.74464 | 0.11231 | 193.60858 | 93.59330 | 87.81537 | 0.13118 | 1.40950 |
| Model_18 | 92 | 0.02187 | 0.04942 | 0.04580 | 0.77032 | 0.11775 | 196.43985 | 90.41224 | 94.96207 | 0.12028 | 1.65727 |
| Model_19 | 77 | 0.03053 | 0.09353 | 0.04637 | 0.78177 | 0.08895 | 162.28150 | 59.24646 | 94.75710 | 0.10751 | 1.37210 |
| Model_20 | 97 | 0.02257 | 0.04722 | 0.03849 | 0.73667 | 0.12359 | 208.20779 | 95.43074 | 99.38693 | 0.13804 | 1.87497 |
| Model_21 | 96 | 0.02223 | 0.04538 | 0.03728 | 0.73107 | 0.12305 | 206.13504 | 95.15987 | 97.58402 | 0.13949 | 1.79067 |
| model_22 | 103 | 0.01558 | 0.02943 | 0.02647 | 0.72129 | 0.13041 | 217.50085 | 97.79645 | 103.47419 | 0.15758 | 1.55107 |
| model_23 | 89 | 0.01731 | 0.02943 | 0.04422 | 0.76497 | 0.09356 | 182.94112 | 97.79645 | 75.17265 | 0.11205 | 1.13231 |
| model_24 | 98 | 0.01589 | 0.02943 | 0.03092 | 0.73670 | 0.11549 | 204.48590 | 97.79645 | 93.06699 | 0.13901 | 1.37754 |
| model_25 | 104 | 0.01507 | 0.02943 | 0.02576 | 0.72694 | 0.12135 | 219.13045 | 97.79645 | 105.72871 | 0.15005 | 1.44667 |
| model_26 | 99 | 0.01649 | 0.02943 | 0.03050 | 0.72485 | 0.13143 | 209.14737 | 97.79645 | 95.56694 | 0.15943 | 1.58594 |
| model_27 | 95 | 0.01678 | 0.02943 | 0.03573 | 0.74248 | 0.10910 | 197.88592 | 97.79645 | 87.81537 | 0.12920 | 1.37825 |
| model_28 | 94 | 0.01657 | 0.02943 | 0.03597 | 0.74664 | 0.11045 | 195.42572 | 97.79645 | 85.13555 | 0.13291 | 1.35494 |
| Model_29 | 84 | 0.02463 | 0.04339 | 0.07254 | 0.78754 | 0.08124 | 176.05014 | 96.80698 | 71.67656 | 0.09008 | 1.23028 |
| model_30 | 104 | 0.01535 | 0.02918 | 0.02562 | 0.71989 | 0.12911 | 219.25808 | 97.33476 | 105.44150 | 0.15848 | 1.57282 |
| Model_31 | 95 | 0.02343 | 0.04634 | 0.04071 | 0.73084 | 0.12706 | 204.76796 | 92.27007 | 98.35500 | 0.14887 | 1.81713 |
| model_32 | 103 | 0.01545 | 0.02918 | 0.02647 | 0.72237 | 0.12659 | 216.13068 | 97.33476 | 103.47419 | 0.14876 | 1.53045 |
| Model_33 | 87 | 0.02636 | 0.05903 | 0.04627 | 0.74967 | 0.12030 | 186.61614 | 82.51329 | 92.84436 | 0.12941 | 1.68782 |
| Model_34 | 95 | 0.02195 | 0.04253 | 0.04597 | 0.75198 | 0.11014 | 203.07412 | 99.10186 | 92.26361 | 0.12325 | 1.70616 |
| model_35 | 96 | 0.01800 | 0.03208 | 0.03024 | 0.71117 | 0.13127 | 201.48598 | 91.04770 | 95.56254 | 0.15496 | 1.53097 |
| model_36 | 96 | 0.01675 | 0.02879 | 0.03394 | 0.73298 | 0.12163 | 201.26120 | 96.98248 | 89.53297 | 0.15360 | 1.48358 |
| model_37 | 89 | 0.02015 | 0.03523 | 0.03802 | 0.72491 | 0.12907 | 186.99572 | 89.97884 | 83.66559 | 0.15001 | 1.57696 |
| model_38 | 88 | 0.01647 | 0.03399 | 0.04022 | 0.77806 | 0.08287 | 180.03567 | 89.56803 | 81.96482 | 0.09662 | 1.02095 |
| model_39 | 80 | 0.02169 | 0.04834 | 0.03802 | 0.74884 | 0.11468 | 165.97922 | 71.03047 | 83.66559 | 0.14104 | 1.49001 |
| model_40 | 96 | 0.01771 | 0.03208 | 0.02983 | 0.71394 | 0.13201 | 201.03122 | 91.04770 | 95.04417 | 0.15562 | 1.54025 |

**Table SI_2: Relevant data to calculate the specific difference of energy ($\Delta E_{SP}$), average participation coefficient ($P_{av}$), Kullback-Leibler divergence (KL), and, relative information centrality ($I_r$) for all possible dimers.**
